# Tail-Robust Quantile Normalization

**DOI:** 10.1101/2020.04.17.046227

**Authors:** Eva Brombacher, Ariane Schad, Clemens Kreutz

## Abstract

High-throughput biological data – such as mass spectrometry-based proteomics data – suffer from systematic non-biological variance, which is introduced by systematic errors such as batch effects. This hinders the estimation of ‘real’ biological signals and, thus, decreases the power of statistical tests and biases the identification of differentially expressed sample classes. To remove such unintended variation, while retaining the biological signal of interest, the analysis workflows for mass spectrometry-based quantification typically comprises normalization steps prior to the statistical analysis of the data. Several normalization methods, such as quantile normalization, have originally been developed for microarray data. However, unlike microarray data, proteomics data may contain features, in the form of protein intensities, that are consistently highly abundant across experimental conditions and, hence, are encountered in the tails of the protein intensity distribution. If such proteins are present, statistical inferences of the intensity profiles of the normalized features are impeded through the increased number of false positive findings due to the biased estimation of the variance of the data. Thus, we developed a, freely available, novel approach: ‘tail-robust quantile normalization’. It extends the traditional quantile normalization to preserve the biological signals of features in the tails of the distribution over experimental conditions and to account for sample-dependent missing values.

---

High-throughput omics data such as mass spectrometry-based proteomics data are subject to systematic non-biological errors such as batch effects, which bias expression level analyses, e.g. in the context of establishing diagnostic and prognostic signatures [1]. Normalization helps in reducing such variations and extracting the relevant biological signal as the main source of the variability that is linked to the biological factor of interest. It oftentimes represents the first data processing step and is critical in quantitative experiments [2], where the choice of the normalization method might influence the downstream analysis results [3] and lead to normalization bias. There exist a multitude of normalization techniques that can be applied in the course of the pre-processing of omics – and in particular proteomics – data. One of these being quantile normalization (QN) [4,5], which, initially, has been applied to microarrays, and was later adopted for proteomics data. It is based on the assumption that at the global level of the whole proteome, the distribution of the protein abundances is similar for all samples. Thus, all quantiles of the measured intensities for each sample are set to the average quantiles over all samples.

QN includes the following steps (assuming it is applied on a matrix where rows correspond to proteins, columns to samples and the values in the matrix cells to intensity values):

1. Sorting the proteins of each sample (column) separately
2. Calculating the mean across each quantile (row) and assigning the mean value to each element in this row
3. Rearranging to the original order of the values in each column.

Yet, as already noted by Bolstad et al., QN proves to be problematic if individual gene expression features occur mainly in the tails of the intensity distributions [5]. Since proteomics data typically comprise features that are in the tails of the distributions, this prerequisite constitutes a serious and practically relevant limitation. Such features demonstrate a small or even no inter-sample variance of ranks across all samples. Depending on the degree of rank invariance they are termed ‘nearly rank-invariant’ (NRI) or ‘rank-invariant’ (RI). A high proportion of rank invariance impedes statistical inferences due to the biased biological variance of the normalized features, which in the case of RI features even results in the latter acquiring the same value across all experimental samples. In addition, it has been found that QN introduces extra patterns into the data on high intensities [6].

Moreover, the abundance of missing values due to technical or biological reasons is a prominent characteristic of proteomics data and can bias every normalization procedure. E.g. it can violate the distributional assumptions that are foundations of QN and may also bias the estimation of the offsets in the course of the here presented tail-robust QN (TRQN), if the occurrence of missing values is correlated with the feature intensities. Low intensity levels that are missing result in over-representation of high values of the same protein. This leads to a biased estimation of the offset and, in turn, may enhance inter-sample distribution differences.

In the present work, we demonstrate the prevalence of features in the tails of the intensity distribution on 173 label-free experimental protein datasets entered to the PRIDE data archive between 01/2013 and 12/2018 that have been processed via the software MaxQuant [7,8] beforehand. Datasets were included if they contain LFQ intensities and – to ensure a representative amount of samples over which the degree of RI is calculated – comprise at least 10 samples. Of these datasets only those proteins with at least 10 data points were included in the analysis. This corresponds to a confidence interval of less than 0.2775 in terms of the percentage of rank invariance. To counteract the normalization bias caused by QN we present a novel modification of the classical QN: TRQN. It is implemented in the R package ‘MBQN’, which is freely available at www.bioconductor.org and github.com/arianeschad/MBQN under the GPL-3 license and also offers a separate function to examine the presence of RI features in omics data.

For a matrix of protein intensities *y*_*ps*_, with proteins p and samples s, each intensity can be thought as sum

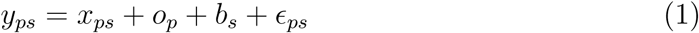

of the ‘true’ biological signal *x*_*ps*_, the protein offset *o*_*p*_, the batch effect of the sample *b*_*s*_ and an error term *ϵ*_*ps*_ ∼ *N* (0, *s*^2^) with sample variance *s*^2^.

TRQN is extending the classical QN by eliminating these offsets *o*_*p*_ for the normalization step. It comprises the following two steps: Prior to the classical QN each feature is transformed to a joint scale by estimating and subtracting the location 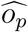 of the distribution of each feature. Then, classical QN is applied. After QN, the normalized intensities are back-transformed to the original scale to preserve the original average intensity levels. Typically, the mean

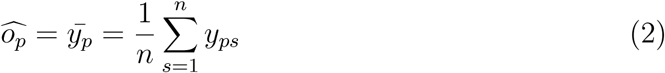

or median can be applied to estimate the location of each feature, although other estimators are also applicable.

Batch effects in proteomics often result in both, decreased intensities and increased number of missing values. In the literature, such missing values are termed MNAR (missing not at random). Figure 1 exemplarily demonstrates the effect of sample-dependencies of MNARs for the PRIDE dataset PXD006138 [9]. For the three highlighted samples with a high number of missing values, the intensity of non-missing features is reduced systematically causing bias in the form of a brighter pattern, or ‘streak’, in heatmaps of the unnormalized feature intensities, which are further aggravated by QN (Figure 1A). The latter enforces equal distribution but introduces a bias in case of sample-dependencies of MNARs. The inter-sample distribution differences are enhanced by MNARs, and, thus, are reduced if only proteins with few missing values are included (compare boxplots in Figure 1B and 1C). By applying TRQN it is possible to attenuate the streaks (Figure 1A).

**Figure 1:**
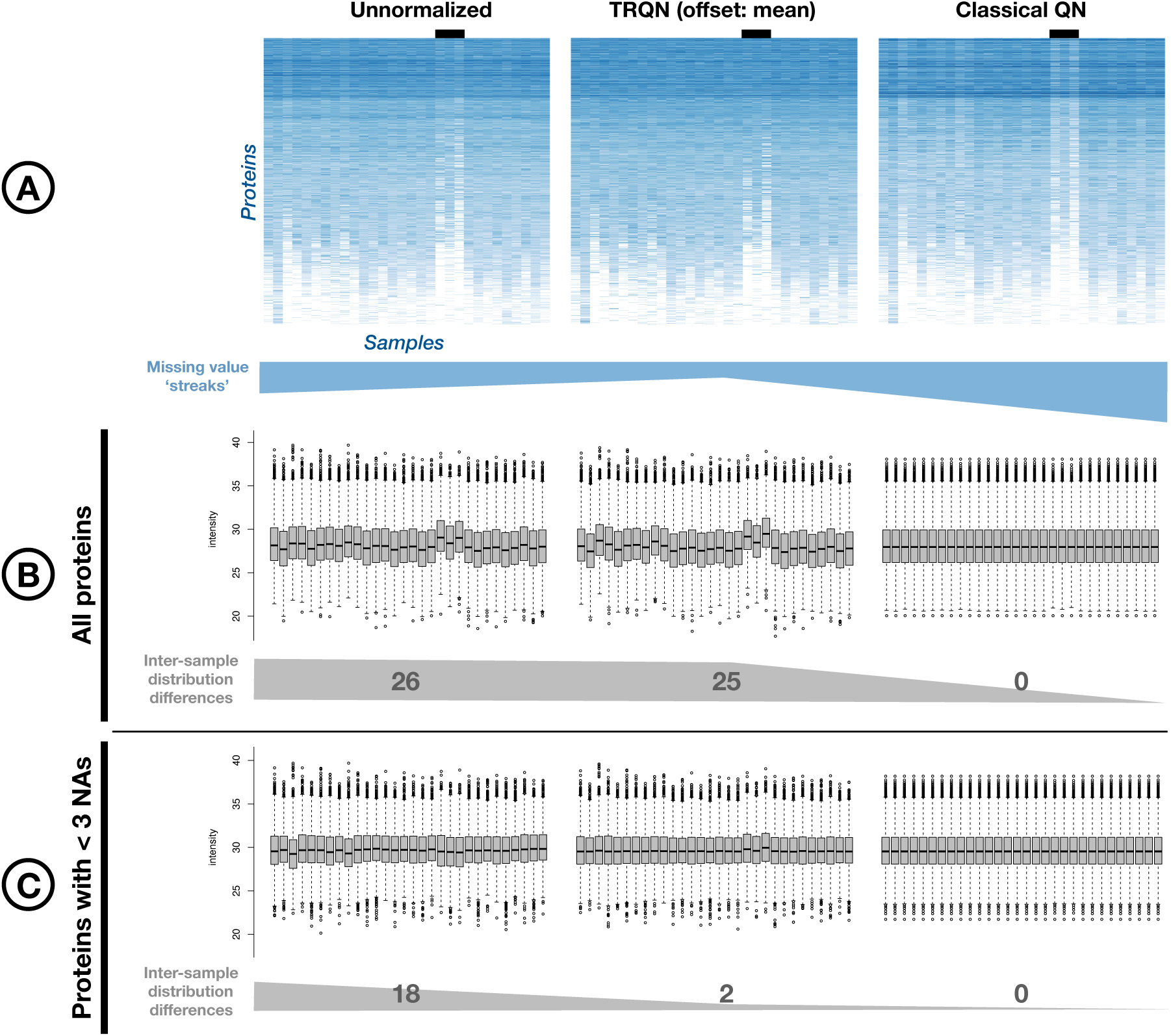
Effect of missing values on inter-sample distribution differences at the example of PXD006138 [9] (30 samples, 9687 proteins, highest RI is 40%) in its unnormalized state and after TRQN with the mean as offset and classical QN. A) Qualitative evaluation of missing value streaks using heatmaps. The protein features (rows) are arranged in descending order of the amount of missing values. The intensity of blue increases with increasing LFQ intensity. The black bars highlight the samples for which these streaks are most prominent. B) and C) visualize the inter-sample distribution differences through the shift in box plots taking into account all proteins and the 3259 proteins with less than three missing values, respectively. As a measure of equality of the distributions of one sample and of the total of the dataset’s intensity values, the number of samples with p<0.05 in the two-sided Kolmogorov-Smirnov test [10,11] is added to both.

**Figure 2:**
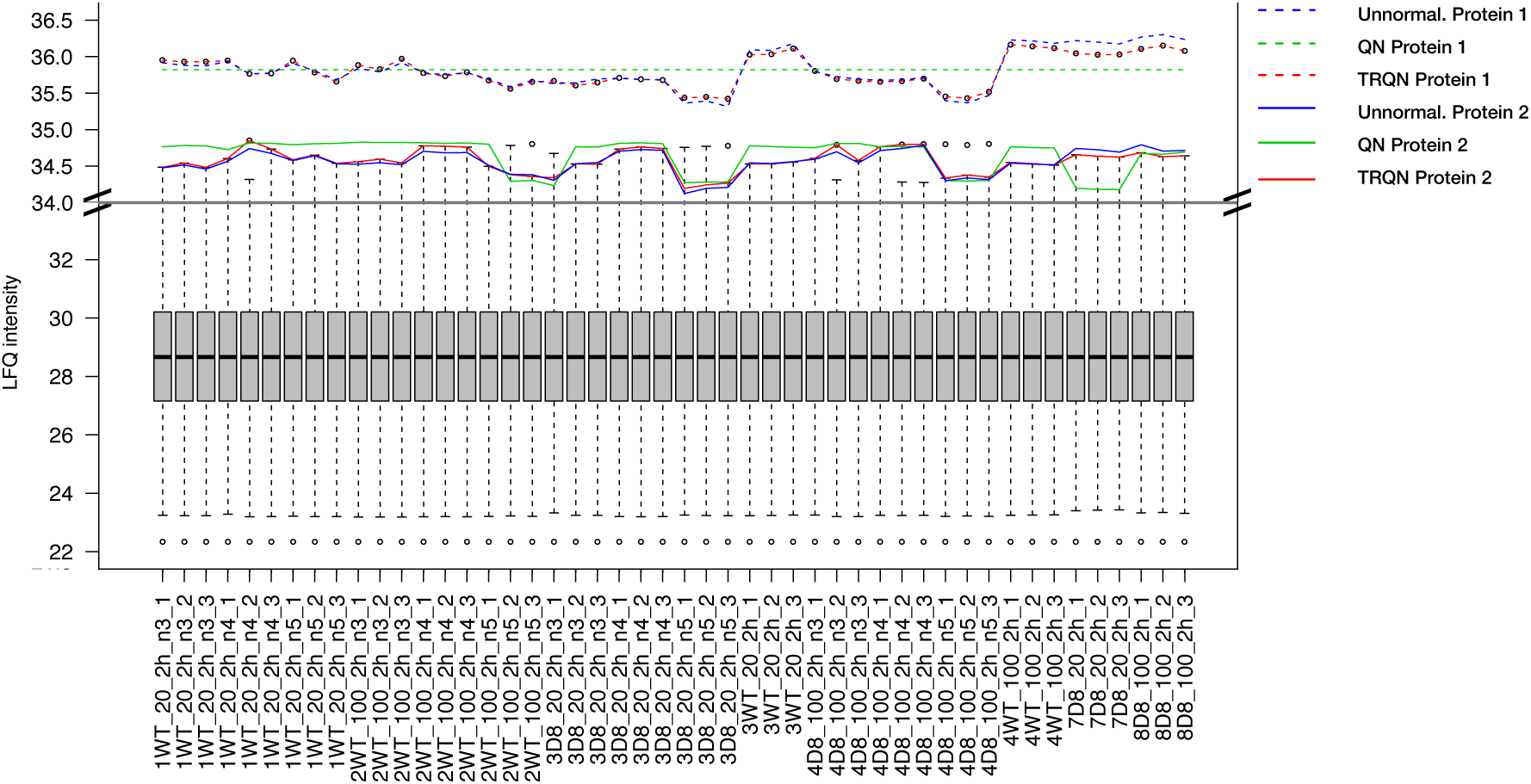
Boxplot of quantile normalized LFQ intensities together with a RI (100 %RI, here referred to as ‘protein 1’) and a NRI feature (75 %RI, here referred to as ‘protein 2’) after QN (green) and TRQN (with mean as offset) (red) across samples. The data are selected from PXD001584 [10] (48 samples, 1223 proteins). The intensity scale above the grey horizontal line is enlarged for a better visualization. Protein 1 (GenInfo Identifier 118498143, elongation factor Tu) is found with vanishing inter-sample variation after QN (dashed green line) that prevents reasonable statistical interpretations. For the nearly rank invariant feature protein 2 (GenInfo Identifier 118498101, glutamate dehydrogenase (NADP+)) QN also leads to almost constant profiles over multiple samples and, thus, to biased variance estimates (green solid line).

As QN has a strong impact on features in the tails leading to the underestimation of the variance, it increases the risk of false positive findings. TRQN, on the other hand, does not have a pronounced impact on proteins in the tails and TRQN normalized profiles are similar to the unnormalized data without such underestimation. Accordingly, we suggest using TRQN instead of QN as it proves to be beneficial both

Finally, 56% (97) of all the 173 investigated PRIDE datasets had at least one protein showing a rank invariance of at least 50%, which would likely benefit from preprocessing with TRQN. A similar picture is likely to emerge from metabolomics data, such as datasets of the MetaboLights database [15]. While our analysis was conducted on the protein level, TRQN can be applied on the peptide level in the same fashion. Also, the advantages of TRQN are not restricted to MS-based proteomics data, but this algorithm may be also useful for other fields of application where data structured in arrays is distorted in scale and location.

## Acknowledgements

We thank our experimental collaborators from the group of Prof. Dr. B. Warscheid, Faculty of Biology, University of Freiburg, Germany, and Prof. Dr. U. Klingmüller, DKFZ Heidelberg, Germany, for helpful discussions.

